# Pupil size and EEG speech tracking as independent measures of listening effort and speech intelligibility

**DOI:** 10.1101/2023.07.31.551390

**Authors:** Ivan Iotzov, Lucas C Parra

## Abstract

Speech is hard to understand when there is background noise. Speech intelligibility and listening effort both affect our ability to understand speech, but the relative contribution of these subjective factors is hard to disentangle. Previous studies suggest that speech intelligibility could be assessed with EEG speech tracking and listening effort via pupil size. However, these measures may be confounded, because poor intelligibility may require a larger effort. To address this we developed a novel word-detection paradigm that allows for a rapid behavioral assessment of speech processing. In this paradigm words appear on the screen during continuous speech similar to closed captioning. In two listening experiments with a total of 51 participants we manipulated intelligibility with changing auditory noise levels and modulated effort by varying monetary reward. Increasing signal-to-noise ratio (SNR) improved detection performance along with EEG speech tracking, suggesting improved intelligibility. Additionally, we find larger pupil size with increased SNR, suggestive of increased effort. Surprisingly, when we modulated both reward and SNR, we found that reward modulated only pupil size, while SNR modulated only EEG speech tracking. We suggest that this new paradigm can be used to independently and objectively assess speech intelligibility and listening effort.

## Introduction

The role of attention in auditory perception is relatively well-studied.[1]–[3] However, the effects of listening effort in the presence of noise and the interaction with attention is not well established.

There are a number of effective techniques for measuring attention during listening tasks, including various measures of ‘neural speech tracking’, which captures the strength of correlation between brain signals and incoming audio.[4]–[8] This phenomenon of an entrainment of neural activity to natural continuous speech has been an active area of study in recent years [7], [9] and shows significant promise as a metric of attention.[10] Speech tracking may also serve as a metric of intelligibility because intelligibility, [1], [11] likely because intelligibility affects the level of attention to the speech. However, there is a lack of clarity as to how these neural tracking measures are affected by other relevant cognitive factors.

Specifically, listening effort is a relevant factor for understanding speech, and it is an important factor when designing hearing aids algorithms. Hearing loss is associated with increased mental fatigue,[12], [13] likely due to the increased effort required to understand speech. While hearing aids may improve intelligibility, fatigue is clearly an important consideration in the design of the devices. [14]. If objective measurement of listening effort were available, it would make a promising target for hearing aid algorithms to reduce fatigue.

Listening effort can be roughly defined as the amount of cognitive resources that are recruited for completion of a task[15]–[17]. There is a long history of using pupil size as a measure of cognitive load or mental activity[18]–[20]. Previous studies have additionally shown that pupil size is a correlate of listening effort[16], [17], [21], with the size of the pupil increasing as effort increases. The pupil is also correlated with changes in attention during tasks[22], [23]. However, the interaction of effort and attention, measured through the pupil size and neural tracking, respectively, is not clear. An additional complication is that pupil size is thought to be regulated by various brain regions (the pretectal olivary nucleus [PON], the superior colliculus [SC], and the locus coeruleus [LC])[20], each of which are involved in diverse cognitive and autonomic functions. Particularly relevant to our study is the role of the locus coeruleus in regulating arousal (i.e. the level of alertness, propensity for action) through the norepinephrine system[20]. It is also well established that initiating movement activates LC and results in phasic pupil dilations[24], [25]. Distinguishing between the tonic activity of LC (baseline level of arousal) and phasic activation due to task demands is difficult and the distinction likely relies on the attention regulation functions of SC, which in turn can influence pupil diameter[26], [27]. It is also useful here to distinguish between pupil size, meaning the diameter of the pupil at a given time, and pupil dilation, which refers to the phasic increase in pupil size associated with task performance or stimulus presentation [17]. The two are often used interchangeably in the literature, but we feel it is important to emphasize the distinction in our case as we are more interested in the tonic pupil size during the task, rather than a phasic dilation in response to a discrete event.

To disentangle the possible links between speech intelligibility and listening effort, we developed a novel paradigm for word detection in noisy conditions that allows for frequent behavioral readout, with the additional advantage of using naturalistic speech. Our approach allows for testing speech detection performance with finer time resolution than typical approaches, in addition to allowing the manipulation of listening effort by varying rewards, and intelligibility by manipulating the speech stimulus. Subjects are presented with continuous natural speech of a narrator telling a story, with a certain level of masking noise introduced over the speech. A single word appears on the screen, similar to a subtitle. The subject is instructed to respond with a button press when they hear the target word uttered by the speaker. Target words are changed every 5 seconds, with some of them serving as false probes that the speaker does not actually utter during the 5 second interval. An important aspect of the paradigm is the reward received for correct responses in the word recognition task. Previous work has shown that reward is an effective way to increase engagement with tasks, especially in motivating participation in more difficult or effortful tasks [28]–[30]. While the main purpose of the reward is to increase engagement in Experiment A, varying the reward in Experiment B provides a way to potentially manipulate the level of effort exerted throughout the experiment.

Through use of this paradigm, we replicate previous results on the effects of changing SNR on word detection performance and neural speech tracking. In addition, we demonstrate that listening effort can be measured independently of speech intelligibility through pupillometry when using reward as an additional experimental factor.

## Methods

### Experimental Paradigm

The experimental paradigm consisted of a behavioral word-detection task. The subject is presented with continuous speech mixed with a varying amount of babble noise. The amplitude of noise varies during the experiment to allow for within-subject comparisons of behavioral and physiological data. The subject is instructed to attend to the stimulus as well as the visual target word, which is presented as text on the screen and changes every 5s. This scenario is familiar to all subjects from their experience of reading subtitles during films. The subject is instructed to press a button as soon as they hear the target word uttered by the narrator. A correct response (true positive) is defined as a button response within 1.5s of auditory presentation of the target word. Any response outside of this window is coded as a false alarm. Behavioral performance in detection tasks is best assessed using the F1 score, a common signal detection metric that is largely independent of criterion shifts. The F1 score is a common metric in signal detection theory and medical imaging [31] and is defined as:

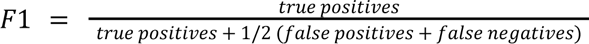

This metric takes into account the detection rate of the subject (i.e. correct responses vs. all possible correct responses), as well as their accuracy (i.e. how many of the button presses were correct vs. how many errors). In this paradigm, true negatives are impossible to measure behaviorally as every word uttered that is not the target word is a true negative. The 1.5s detection window was chosen *a priori* based on previous work [1]. In order to verify that this was a reasonable choice of reaction time window, we calculated the F1 score across all subjects using a variety of window sizes (see Fig. 3E). Indeed, we do see an inflection point around 1.5s, demonstrating the validity of the window choice.

Additionally, the two experiments had different reward paradigms. In Experiment A, participants were rewarded for each correct response across the whole experiment. This constant reward was included in order to maintain a steady level of engagement throughout the task and ensure the participants’ attention. In Experiment B, two different reward conditions were used. Participants were either rewarded for each correct response in one condition, or they were given no reward in the other. This manipulation was intended to modulate the level of effort exerted by the participants in each condition, with high effort in the rewarded condition and low effort in the no-reward condition.

### Stimulus Description & Presentation

All stimuli in these experiments were presented using Sony MDR 7506 headphones at ∼70dB SPL in an otherwise quiet environment. Volume level was calibrated by measuring the output of the headphones using a ExTech Instruments 407730 sound level meter fitted with a rubber coupling meant to simulate the ear canal. Two different stimuli were presented to subjects in two different experiments that all used the experimental paradigm described above:

Experiment A used an audio narrative with a run-time of 13 minutes, namely, Chapter 1 of *The Name of the Wind* by Patrick Rothfuss. The audiobook is professionally narrated by a male speaker with a standard American accent. Target words are displayed on the screen every 5 seconds. Subjects in this experiment were given a fixed monetary reward for each word they correctly identified ($0.25 per correct detection), in addition to a base fee to compensate them for their participation ($30). The average compensation received was $55, with a maximum of $70 and a minimum of $40. The stimulus was divided into 8 sections, switching in SNR between −9dB and −3dB of additive babble noise. This resulted in a total of N=37 words per SNR condition.

Experiment B used an audio narrative with a run time of 30 minutes, with Chapter 2 of *The Name of the Wind*, narrated by the same individual as in Experiment A. This stimulus is divided into 8 sections of 224.5s each in order to allow subjects to rest and to prevent fatigue during the task. The experiment consisted of 4 conditions, 2 different levels of babble noise (−9dB and −3dB SNR, interleaved as before), and 2 different levels of reward (reward of $0.20 per correct detection, and no reward or instant feedback). Similar to Experiment A, subjects were rewarded for each word that was correctly identified in the reward blocks, but received no feedback on their responses in the no-reward block. The average compensation received was $51, with a maximum of $70 and a minimum of $45. The subjects were alerted before the start of each block whether they would receive rewards for that block, or not. The order of conditions was randomized between subjects. Target words were again presented every 5 seconds resulting in a total of N=45 words per noise/reward condition.

For both experiments, target words were selected to contain a mix of different phonetic and semantic properties across conditions. Functions words (e.g. ‘the’, ‘and’, ‘to’) were excluded from the possible choices for target words, along with most proper nouns except for the names of the main characters. Though the task is a word detection task, the surrounding context for each word enables participants to more easily predict and detect the target word. This is intentional, as it transforms the task into a more generally intelligibility focused one, rather than simple word detection.

### Subjects

All experiments were conducted at the City College of New York (CCNY). Subjects were recruited through advertisements posted around CCNY campus. Subjects were screened for self-reported normal hearing. All recruitment material was approved by the CUNY Institutional Review Board. 31 subjects were recruited for Experiment A (18 female, age range = 18 - 41, mean age = 21.8 ± 5.3), and 20 for Experiment B (15 female, age range = 18 - 39, mean age = 22.5 ± 5.6). The sample sizes for each experiment were chosen based on previous work[1] which achieved robust results with a sample size of 20. We kept the same sample size for Experiment B, but due to Experiment A being significantly shorter, elected to use 35 subjects for Experiment A. After collection, data from 4 participants were excluded due to poor data quality, either with significant EEG recording issues or missing 10% or more of the pupil data.

### EEG Recording & Preprocessing

EEG recordings were performed using a BioSemi Active II amplifier with 64 electrodes arranged according to International 10-20 standards. Additionally, 6 electrooculogram (EOG) electrodes were applied, 3 arranged around each eye. EEG was sampled at 1024 Hz and saved using BioSemi ActiView software. Presentation of the stimulus was done using MATLAB and Psychtoolbox software[32].

Offline, EEG data was filtered using a 5th order high-pass Butterworth filter with a cutoff of 0.5Hz in order to remove low frequency fluctuations. Eye movement artifacts were removed by regressing out the EOG signals from the EEG using least-squares linear regression (see <u>parralab.org/isc/</u>). Outlier values more than 3 standard deviations from the mean in each channel were identified and set to 0, along with 20ms preceding and following the outlier. This criteria was chosen because ∼99.7% of the data is contained within 3 standard deviations of the mean, so values falling outside of that range are clearly extreme. Finally, EEG data were downsampled to 128Hz to reduce processing time.

### Pupil Recordings

Eye tracking data was collected for Experiments A and B. Luminance was controlled during the experiments, and target words were presented in color font that were equal in luminance to the gray background. The relative luminance of a color can be found using the sRGB definitions of the relative luminance of each primary color. An equal luminance color for the text presented was then selected based on the relative luminance of the background. This process ensures that the pupil does not react to changes in the target word on the screen. Pilot experiments confirmed that visual word presentation did not elicit a pupil response and this is visualized in Fig. 3D.

For Experiment A, eye tracking data was collected using an SR Research EyeLink 1000 Plus camera recording at 512 Hz. The tracking camera was calibrated for each subject before the start of the experiment using SR Research software and recorded using the same.

For Experiment B, eye tracking data was collected using a Tobii Pro Fusion camera recording at 250 Hz. Calibration was done using Tobii Pro Eye Tracker Manager and recorded using MATLAB.

Before the start of the experiment, subjects were asked to sit quietly for 5 minutes to establish a baseline before stimulus presentation. The mean size of the pupil over this rest period served as the baseline pupil size.

### EEG Stimulus-Response Correlation (SRC)

In much of the auditory attention literature, attention is estimated using a forward or backward model[3], [33]. In the forward model, the amplitude envelope of the clean speech, *s(i)*, is projected using a set of learned filters and then correlated with the brain response in each EEG channel, *r(i)*. This essentially models the brain response as a linear encoding of the input stimulus. The backward model is the opposite, where the brain response is used to reconstruct the input stimulus and this reconstruction is correlated with the original stimulus. The model used here is a hybrid of these two approaches. The model learns a set of temporal filters for the stimulus (Fig. 1B) and spatial weights for the neural signals (Fig. 1A) so as to maximize the correlation between the projected signals using canonical correlation analysis[34]. The final stimulus-response correlation (SRC) value is calculated by projecting all the data for a given subject and condition using the learned filters and weights and calculating the Pearson correlation between the filtered stimulus and projected EEG response. For a more complete description of the SRC method, please see (Dmochowski et al.)[35].

**Figure 1.**
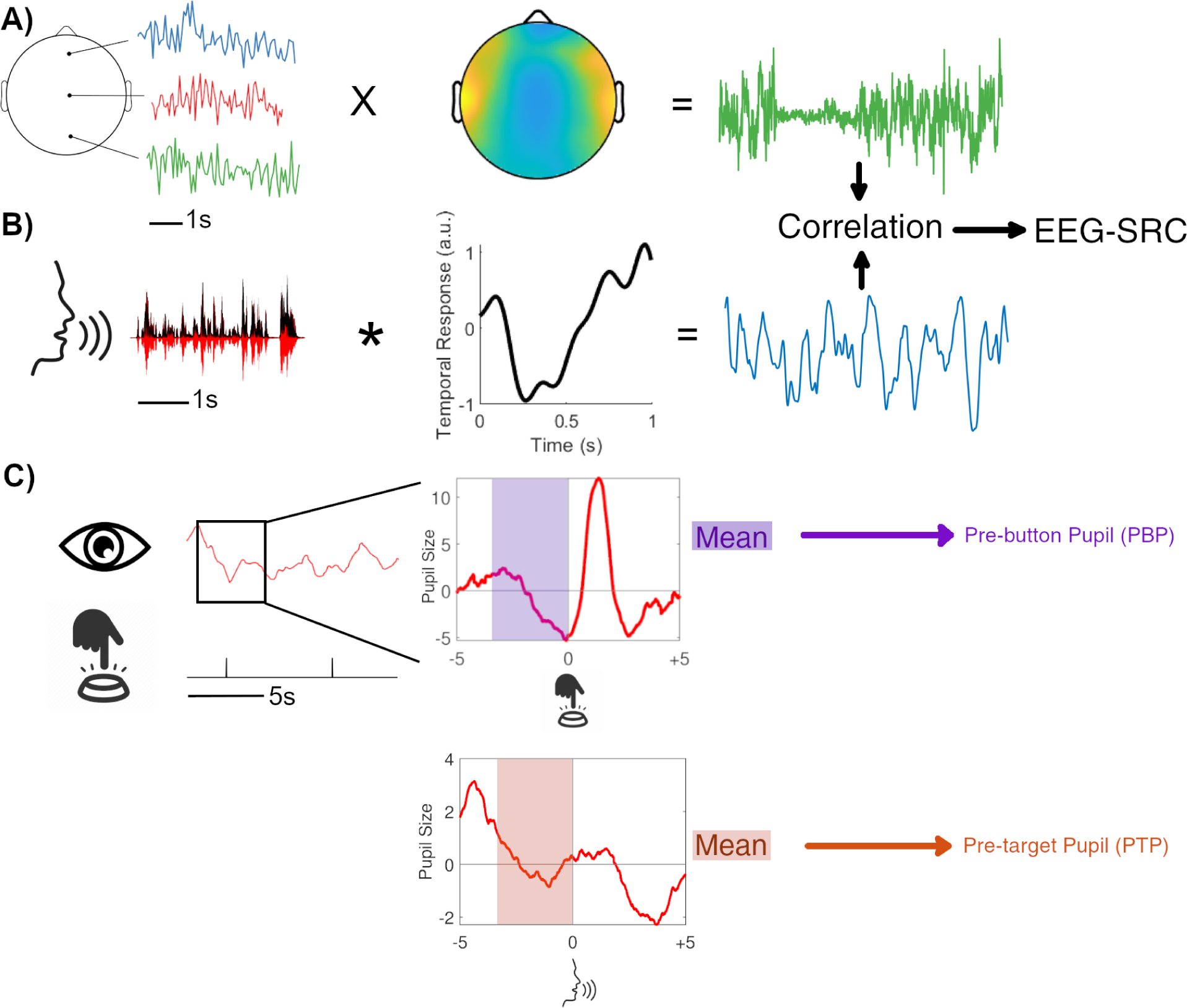
Illustration of various metrics used throughout this paper. The stimulus-response correlation (SRC) is calculated by projecting the EEG responses (A) using a bank of spatial weights, projecting the stimulus (B) using a bank of temporal filters, and then correlating the two to produce a final EEG-SRC score for each condition. The pupil size metrics used are shown in (C). In purple is the pre-button pupil size (PBP), the mean pupil size in the 3s preceding a button response as % of baseline pupil size. In orange is the pre-target pupil size (PTP), the average pupil size over the 3s preceding an auditory target word presentation.

A more traditional approach would be to use a temporal response function (TRF) or similar to quantify the response. However, there are two advantages to our approach over the TRF. First, we are not interested in the response at specific electrode locations and wish to avoid the additional complication that is presented by electrode selection and comparison. Second, our paradigm is a continuous auditory stimulus, so choosing a specific time point at which to align the TRF would be somewhat arbitrary. Moreover, we are not interested in the instantaneous response to, for example, target word presentation. Instead, we examine how well-represented the stimulus is in the brain response across conditions, which the SRC method is better suited for.

### Pupil Preprocessing & Analysis

For both experiments, after calibration of the eye tracker, subjects were instructed to keep their gaze focused on the screen and sit still as the experiment was carried out. Offline, blinks and other outliers were identified based on the interquartile range and removed. Then, these gaps in the signal were filled using linear interpolation. Additionally, the signals were filtered using a 250 ms median filter.

Finally, all pupil data were downsampled to 128Hz. Pupil data were normalized for each subject and are reported here as a percentage of the baseline pupil size. This transformation allows for comparison between subjects and between the two different types of eye trackers. Because the baseline is taken in a period of rest before the beginning of the experimental task, all pupil size values reported are likely to be higher than the baseline due to participants’ engagement in the task itself.

We examine two different pupil metrics, each aligned to a different moment in the detection task, and used for different purposes (Fig. 1C). The primary metric is the mean of the pupil size in the 3s before the spoken target word, regardless of whether the word was detected or not. This metric is averaged across all words in a condition and is referred to as pre-target pupil size (PTP). Its primary purpose is to capture the state of the listener in different noise and reward conditions, unaffected by word detection or button response. The second metric is the pupil size in the 3s preceding a button response, regardless of whether it was a true or false positive. Unlike the PTP, here we analyze results on a single trial basis (i.e. each button press is treated as a separate trial). The purpose of this metric is to compare the state of the listener prior to correct and incorrect button responses. This second metric will be referred to as the pre-button pupil size (PBP). In the case of PTP, we are interested in comparing the state of the participant across the various conditions. In the case of PBP, we are interested in linking pupil size to behavioral performance on a single trial basis. Due to the nature of the experimental task, two different pupil measures are necessary in order to capture every possible response (true positive, false positive, and false negative). False positives cannot be captured by the PTP because there is not an auditory target present that can be responded to. Similarly, false negatives cannot be captured by PBP because the measure necessitates a button response. Therefore, both measures are necessary in order to analyze the full range of behavior. Finally, we use a Bayes factor analysis in Experiment B to determine the strength of evidence for the null results [36]. The bayesFactor toolbox [37] was used for the calculations, and the default scale of for the alternate hypothesis was used to calculate the Bayes 2/2 factor for the ANOVA results.

## Results

### Effects of Noise on Neural Tracking and Pupil size (Experiment A)

The purpose of Experiment A was to validate the use of the new word detection paradigm as well as study the effects of masking noise on pupil response in this context. In order to increase engagement with the task, participants received immediate feedback of a reward for each target word they correctly identified, and reward for each correct word was held constant. As expected, word detection performance as measured by the F1 score was significantly higher in the −3dB than in the −9dB condition (Fig. 2A, paired sample t-test, t(30) = 9.72, p = 8.7×10^−11^]. This finding is reinforced by reaction time data (Fig. 3C), with reaction time being significantly faster in the −3dB condition compared to the −9dB (Wilcoxon rank sum test [Z = −6.8, p = 9.1 x 10-12]).

**Figure 2.**
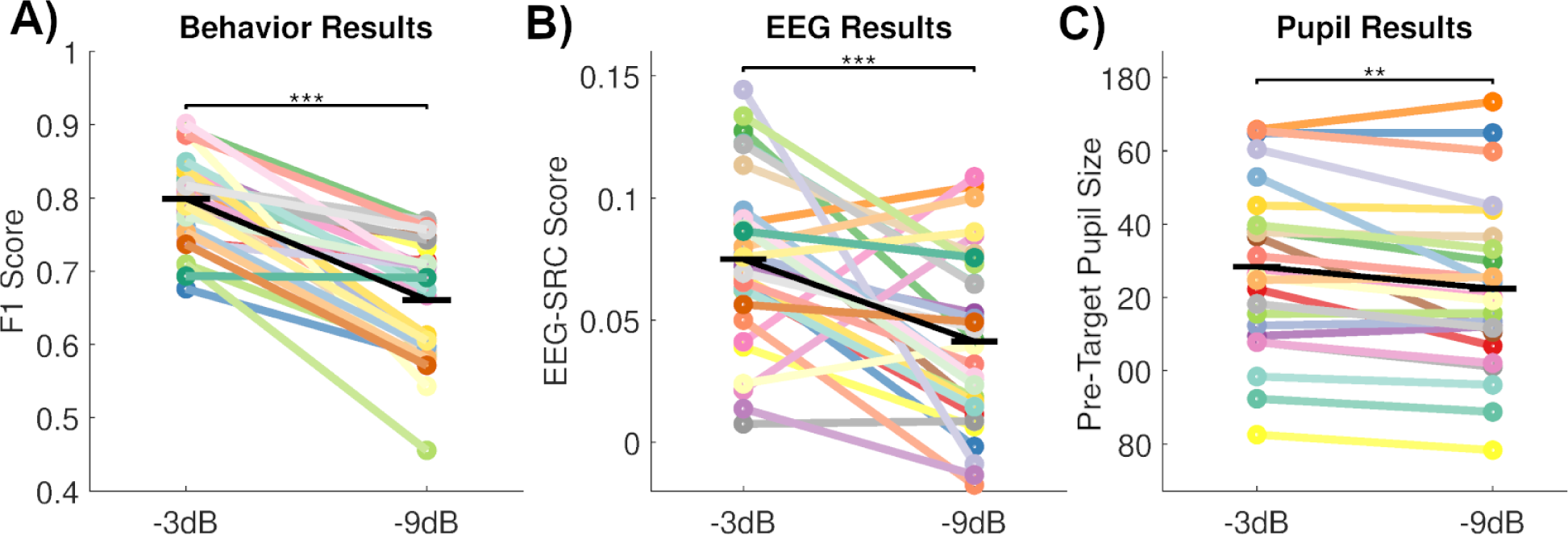
Behavioral, EEG and pupil results of Experiment A. (A) shows behavioral results for each condition in terms of F1 score. (B) shows the EEG-SRC between the clean speech stimulus and EEG responses for each condition. (C) shows the pre-target pupil size results for each condition. *, ** and ** indicate p-values of 0.05, 0.01 and 0.001 in the mixed effect ANOVA in the main text (N = 31).

In order to evaluate the effect of noise on the brain, we compare the EEG-SRC within subjects between the two SNR conditions (Fig. 2B). This metric captures how correlated the EEG signals are to the envelope of the clean speech. A 2-way mixed-model ANOVA with noise level as a fixed effect, and subject identity as a random effect shows a significant effect for noise level [F(1, 30) = 17.73, p = 2×10^−4^]. These results fall in line with previous work showing increased EEG speech tracking with improved SNR [11], [38].

To determine the effect of noise condition on pupil size, we first examined the mean pupil size 3s before auditory target word presentation (PTP, see Methods) averaged over all target words within SNR condition (Fig. 2C). We applied a two-way mixed model ANOVA on these data, with a fixed effect of noise level and a random effect of subject identity. We see a smaller, but significant increase of PTP with improved SNR [F(1,1799) = 12.83, p = 0.002]. This analysis indicates that the task demands were sufficient to elicit differentiable pupil responses. However, this result runs somewhat contrary to that typically found in the literature; increasing pupil size with increasing task demands. This could be a reflection of effort allocation strategies on the part of the subjects. Previous work suggests that listeners prioritize the rewards that are relatively easier to realize in the context of a rewarded task [39].

To relate pupil size directly to behavioral responses we also analyzed on a single-trial basis the pupil size 3s prior to each button response (PBP, see Methods). We applied a two-way ANOVA to PBP, with noise level and result of the button press (i.e. true positive or false positive) as fixed effects. Again we find a significantly larger pupil in the higher SNR condition [F(1,1624) = 8.44, p = 0.0037], as well as a larger pupil for incorrect responses [F(1,1624) = 6.28, p = 0.0123], i.e. pupil size is larger preceding false positive responses as compared to true positives (Fig. 3B). This result is consistent with an increased level of arousal (as measured by pupil size[40]–[42]) lowering participants’ response thresholds, leading to an increase in false alarms.

**Figure 3.**
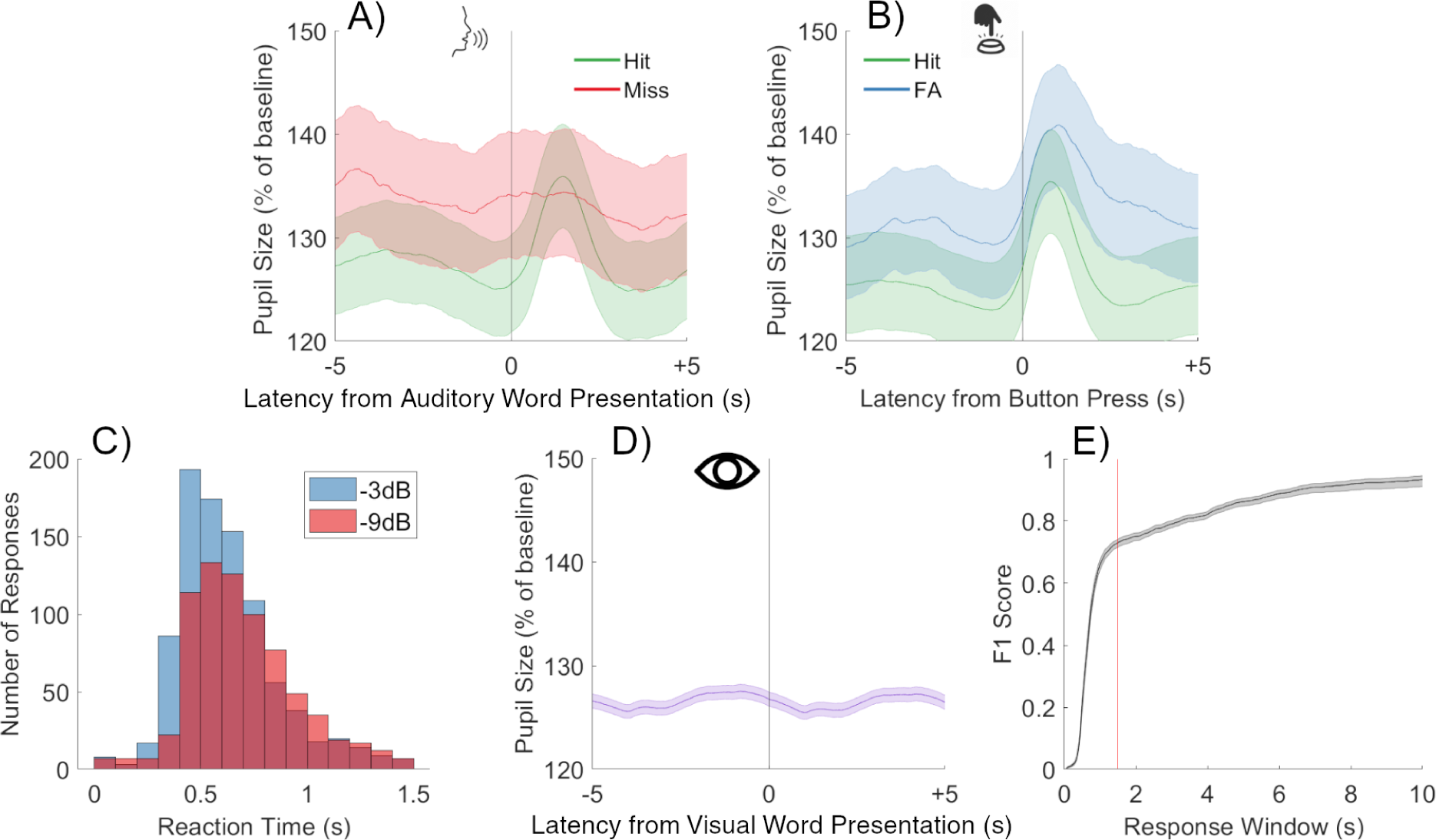
Pupil and behavior as a function of time. (A) Pupil size averaged over all trials in which the subject correctly identified the target word compared with all trials in which the word was not detected. Time = 0 corresponds to the time the target word is uttered by the speaker. Shaded areas represent the standard error (N = 31) (B) Average pupil size over all trials in which the target word was detected, compared with false positive trials. Time = 0 corresponds to the time the button was pressed. (C) Reaction time to correctly identify target words for the two SNR conditions. (D) Pupil size aligned to target presentation as text on the screen. (E) Dependence of F1 score (averaged across subjects) as a function of the detection window duration, standard error across subjects is shown in the shaded region. We used 1.5s (red).

In Figs. 3A and 3B, we see an obvious phasic pupil dilation associated with the button response. We measure its size as the mean pupil difference between pupil size 3s before and after the button push. To determine if this phasic dilation is related to the SNR condition or the accuracy of the response we performed a 2-way ANOVA with the effects of noise level and response accuracy. We found no effect of response accuracy on the post-button pupil dilation [F(1,1624) = 0.38, p = 0.54], and no effect for the SNR condition as well [F(1,1624) = 0.03, p = 0.86]. Given these results, we concluded that this post-button response is likely caused by the motor action of the button press. We proceed to use the pre-target and pre-button measures described above, as they are not affected by this button response artifact.

### Effects of Reward and SNR on Pupil Size and EEG SRC (Experiment B)

In Experiment B we intended to modulate effort by varying the level of reward for correct detections. Subjects were presented with the same reward paradigm as in Experiment A, except that in half the trials there was a reward for correct responses and in the other half no reward was offered. The stimulus was divided into 8 equal sections of 4 conditions (−3dB reward, −3dB no reward, −9dB reward, and −9dB no reward). Condition order was randomized between subjects and subjects were informed before the start of each block whether there would be rewards, or not.

Replicating the previous experiment, there was a strong effect found for noise in terms of the F1 behavioral performance (Fig. 4A) [F(1,19) = 218.31, p = 7×10^−12^]. We found a small, but significant improvement in behavioral performance with reward [F(1,19) = 4.92, p = 0.039]. The reaction times of participants were also significantly faster in the −3dB condition (Wilcoxon rank sum test [Z = −11.79, p = 4.2 x 10^−32^]), and when they were rewarded (Wilcoxon rank sum test [Z = −2.0, p = 0.042]).

**Figure 4.**
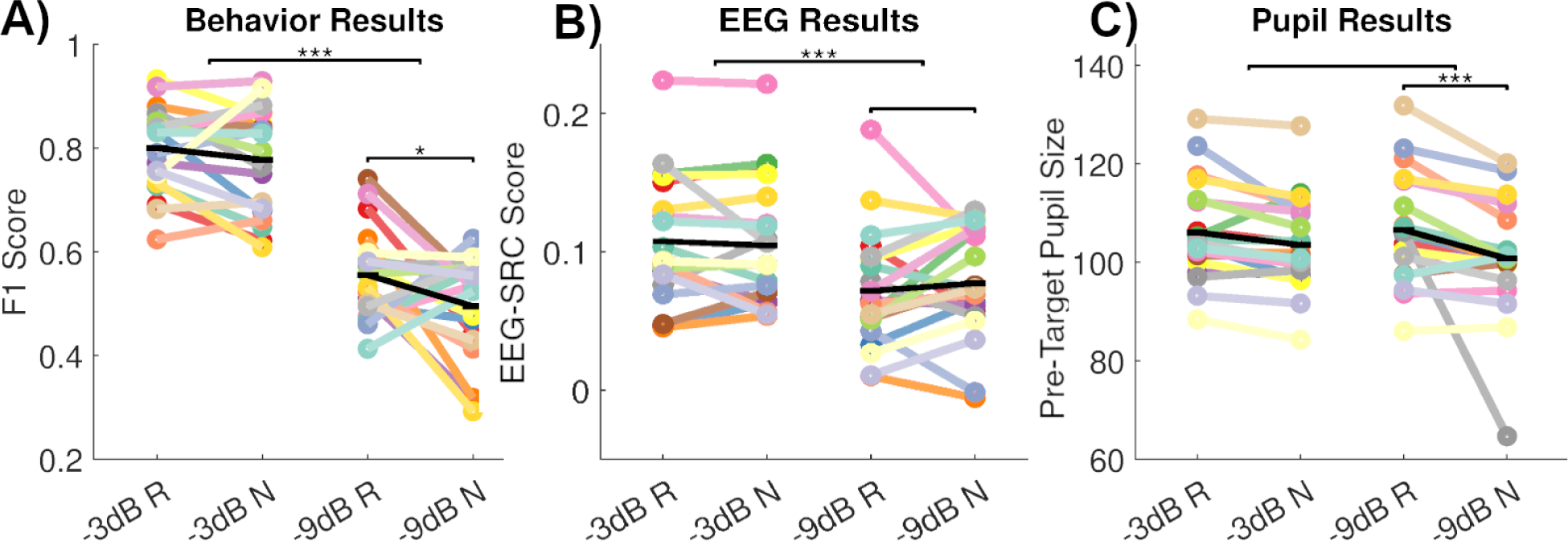
Behavioral, EEG and pupil results of Experiment B. (A) shows behavioral data in terms of F1 score for each of the 4 conditions. Significant effects were found for noise, and a small effect for reward. (B) shows the EEG-SRC results in each condition. No effect was found for the reward condition, but a strong effect for noise. (C) shows the pupil size before target word presentation, as detailed in the Methods and in Fig. 1. Strong effects were found for reward, but not for the noise level. *, ** and ** indicate p-values of 0.05, 0.01 and 0.001 in the mixed effect ANOVA in the main text (N=20).

As in Experiment A, we used SRC analysis to compare the EEG responses between conditions and subjects (Figure 4B). We used a 3-way mixed-model ANOVA, with fixed effects of noise level and reward level and a random effect of subject. We found a strong increase in EEG-SRC with improved noise condition [F(1,19) = 30.20, p = 2.6×10^−5^]. Importantly, there was no significant effect of reward on the EEG responses [F(1,19) = 0.07, p = 0.80]. This data is contrary to our expectations that the changes in effort produced in the subject by the variable reward would create differences in attention, and thus in the EEG SRC metric, known to be modulated by attention to the stimulus[43]–[46]. The results, however, are consistent with the interpretation of EEG-SRC as a marker for intelligibility as EEG-SRC increased with SNR.

We also examined the mean pupil size for the 3s before target word presentation (PTP) averaged over all targets (Fig. 4C). We used a 3-way mixed-effects model ANOVA with the fixed effects of noise and reward level and a random effect of subject identity. We find that in the reward condition pupil size preceding word presentation was larger [F(1,99) = 15.17, p = 0.001], but there was no effect for the level of noise [F(1,99) = 1.43, p = 0.25]. These results seem to indicate that the main driver of the pupillary response is the level of reward offered to the subject. We believe that this difference in reward is driving differences in the participants’ effort levels, and thus the pupil response. But, it seems that modulating reward does not drive the EEG response in the same way as the pupil (See Fig. 5A). In order to quantify the strength of the evidence for a null result in regards to noise and the pupil size, we performed a Bayes factor analysis on the factor of noise. We find a Bayes factor value of 4.0 in regards to the noise null result, indicating strong, but not exceptional, evidence for our findings.

**Figure 5.**
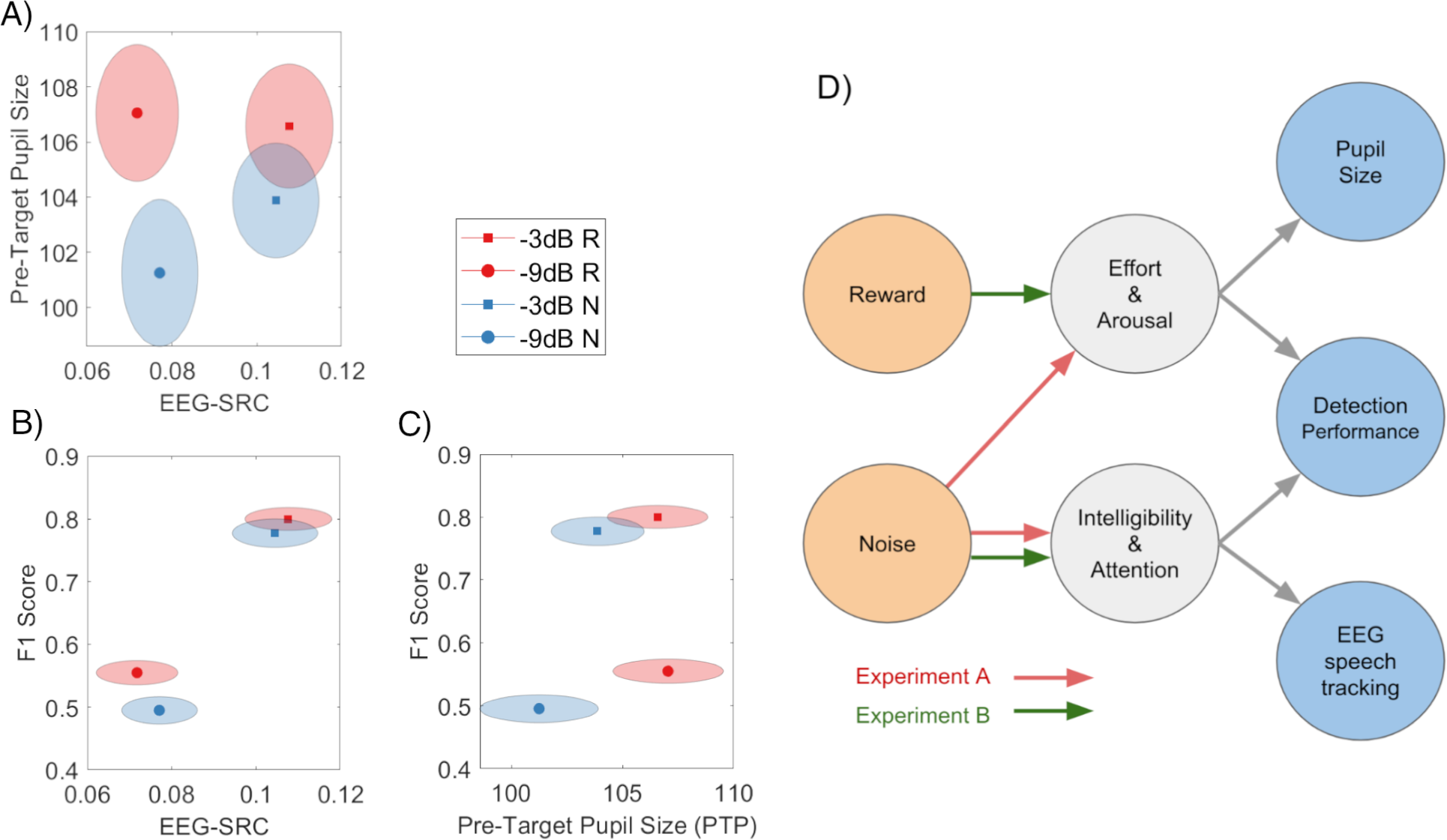
Summary of results and conceptual model. (A) EEG speech tracking (EEG-SRC) vs. pre-target pupil size. Color indicates the reward condition and symbol shape indicates the SNR level. The mean across all participants is plotted, with surrounding shading showing the standard error. (N = 20). (B) behavioral performance (F1) EEG speech tracking (EEG-SRC). (C) Behavioral performance vs pre-target pupil size. (D) Conceptual model of experimental control variables (orange), psychological constructs that are not directly observable (gray), and observable outcome variables (blue). Arrows indicate hypothesized causal relationships consistent with the observed results in Experiments A and B.

We also repeated the single-trial analysis to link pupil size to the behavioral performance within subjects. We looked at the mean pupil size in the 3s preceding button responses (PBP). We evaluated this data with a 4-way ANOVA mixed-effects model with noise level, reward level, and whether the word was later correctly identified as fixed effects, and subject identity as a random effect. As in Experiment A, we find a significantly larger pupil for false alarms [F(1,5177) = 9.63, p = 0.0054] suggestive of an effect of arousal. And again, we found a significantly larger pupil in blocks that provided a reward [F(1,5177) = 10.88, p = 0.0018]. However, unlike in Experiment A, we did not find a significant effect of noise level on the pupil preceding a button push [F(1,5177) = 0.13, p = 0.72]. These results parallel those found for the PTP metric indicating that pupil size captures a state of the subject in the different conditions regardless of whether pupil size is measured prior to the target stimulus or prior to a motor response.

The results of Experiment B are summarized in Fig. 5A-C. The most important observation (Fig. 5A) is that reward seems to affect only pupil size, while SNR affects only EEG speech tracking (EEG-SRC). Behavioral detection performance measured as F1 is affected by both (Fig. 5B and 5C).

## Discussion

We aimed to investigate the effects of listening effort and speech intelligibility on neural tracking and pupil size through manipulation of reward and SNR. We found that decreasing SNR has a negative impact on neural speech tracking measured through SRC, and that increasing reward increases the size of the pupil. Additionally, we found that these two effects can be separated and thus measured independently, given careful experimental setup.

We interpret these results in terms of effort and arousal on the one hand, and intelligibility and attention on the other. These are psychological constructs that, arguably, are hard to quantify objectively. Fig. 5D summarizes our interpretation into a conceptual casual model. In Experiment B, efforts and arousal, which modulate pupils size were affected only by reward, while intelligibility or attention, which modulate EEG-SRC were only affected by SNR. In contrast, in Experiment A, SNR may have affected both intelligibility and effort and thus SNR modulated both physiological metrics. In this view, the particular contribution of this work is to have presented an experimental paradigm that can independently assess listening effort and speech intelligibility with distinct and objective physiological markers for each.

Our design also has certain advantages over previous work investigating the relationship of listening effort and pupil size, which mostly rely on single sentence recall tasks[17], [47], [48]. The advantages of continuous speech over the conventional use of single sentences are threefold: First, behavioral responses are more frequent and the time resolution can be varied based on experimental demands. Second, the stimuli are more naturalistic than single sentences, allowing for investigation into more naturalistic contexts, which may be important in the context of speech tracking, etc. Finally, the stimulus used can be changed easily (we demonstrate the use of 2 different speech stimuli), allowing for presentation of varied stimuli to investigate various aspects of hearing in noise without changing the underlying paradigm.

Some previous work shows that, at least in online scenarios, reward improves engagement with auditory tasks [30]. Other work done in lab scenarios does not show the same kind of improvements in engagement/performance using reward [49], [50], though the authors report differences in effort measured through pupil size as a result of the changing rewards. This can be partially explained through differences in motivation, as online subjects have a different set of motivations and incentives for their participation compared with subjects that come into a lab. Additionally, part of this difference could be explained through participants’ allocation of resources in order to maximize their reward while minimizing the amount of effort exerted [39]. An important aspect of the work presented here is the difference between the incentive structure of the two experiments, one with constant reward and one with variable reward. Having a reward present for both the easier and harder SNR level encourages a strategy of prioritizing the easier SNR, as the level of effort vs. reward is more advantageous in easier circumstances. Varying the reward throughout the experiment presents more limited opportunities to receive the reward, thus changing the incentive structure present in the constant reward scenario.

Note that we measured pupil size preceding stimulus or response with a baseline at the very beginning of the experiment. This differs from a common approach of measuring pupil dilation, where change in pupil size relative to some period immediately before a trial is often measured [17], [50], [51]. The only phasic change in pupil size we saw in the continuous task was associated with the button press, and this dilation was not predictive of task performance, nor did it correlate with any of the task conditions. In this sense, in this work we are not technically measuring pupil “dilation” in the conventional sense. Rather, we measured pupil size preceding the individual stimulus/response and thus are capturing a state of the user that did fluctuate across trials and conditions.

### Effects of Noise

We hypothesized that decreasing signal-to-noise ratio in the stimulus will lead to a decrease in neural tracking as measured through EEE-SRC. Additionally, we predicted that the changes in SNR would have some impact on pupil dilation as listeners have to exert more effort in order to understand in more adverse conditions. As predicted, decreasing SNR did indeed lead to lower levels of neural tracking. This result is in agreement with our previous work in similar paradigms[1]. We believe that this is due to attentional effects, where if the listener is unable to comprehend incoming speech signals, their attention wanes and therefore neural tracking is decreased. Of course, this is not the only relevant factor, as the noisier speech signal in lower SNR stimuli contains degraded speech information and blurred acoustic boundaries, thus making it more difficult to track. Behavioral results from the different noise conditions show a dramatic decrease in performance on the behavioral task with decreasing SNR. However, some recent work[52] has shown that neural tracking may be a necessary condition for intelligibility, but not a sufficient one. In a speech rate paradigm Verschueren et al. showed an increase in neural tracking along with a decrease in behavioral intelligibility measures with increasing speech rate.

These results, along with our own, suggest that the link between intelligibility and neural tracking depends on the exact experimental conditions. By itself, neural tracking does not capture higher-level aspects of speech comprehension tied to linguistic components of the speech signal.

Fluctuations in pupil size due to changes in SNR was also an important measure of interest in our study. We hypothesized that SNR would have a strong effect on the pupil, as indicated by the literature on the subject[16], [17], [48], [53]. Indeed, in the constant reward experiment (Experiment A), we found a significant effect of noise on pupil dilation. However, despite having an identical behavioral task, the variable reward experiment (Experiment B) did not produce significant differences in pupil dilation between the noise conditions.

### Effects of Reward

An additional hypothesis in our study was that offering variable rewards to participants for their performance would affect the effort they exert on the task, and thus affect their pupil size. This hypothesis was borne out by the results, with the participants’ pupil size increasing significantly in the reward condition. This effect was especially pronounced in the noisier condition. We believe this is because the noisier condition is inherently more demanding, thus requiring a larger recruitment of cognitive resources in order to compensate for the more degraded speech. This is supported by the literature, where pupil dilation is increased in conditions where it is more difficult to focus attention to a stimulus[51], particularly in the presence of a competing speaker[47]. Pupil dilation increases in the easier condition as well, but less effort is required in order to increase performance in the less challenging condition.

Interestingly, changes in the noise level in Experiment B did not produce significant changes in the pupil, while noise level did have a significant effect in Experiment A. We believe this is due to the reward structure of each of the experiments[54]. Participants in Experiment A were rewarded for each correct response in both noise conditions. Because the noisier condition is inherently more demanding, they increased their level of effort in the noisier condition in order to compensate for the more degraded speech. However, in Experiment B, the more relevant factor was the level of reward rather than the level of noise. Because there is no benefit in exerting more effort in the unrewarded conditions, regardless of noise level, we see no changes in the pupil due to noise alone.

### Limitations

There are several limitations in the studies presented above. Firstly, while the behavioral task does seem to measure word detection effectively, the task itself is not how everyday listening is done. The presence of the target word on the screen creates a scenario where subjects may be listening specifically for the target word, to the detriment of their overall comprehension. There is no analogue to this in real-world conditions.

Secondly, the button response to indicate identification of a target word may be a confounding factor in the pupil response. Namely, it could be the case that part of the difference in pupil response in the hits vs. misses could be due to the motor signals that are required to press the button. However, this is unlikely as the results on pupil size hold even when pupil size was measured prior to auditory presentation of the target word, i.e. when button press would not have been initiated yet.

Thirdly, there has been considerable disagreement about the role of neural speech tracking in speech comprehension and intelligibility[52]. So it is not clear if the present results on intelligibility are indicative of overall speech comprehension. Future studies may use post-experiment comprehension questions to determine if these results carry over to the practically more relevant issue of speech comprehension.

Finally, our conceptual model is only one of the possible interpretations of the result. For instance, it is quite possible that the intelligibility affected effort. Future work may add post-experiment questionnaires for subjective judgment of effort and intelligibility. We note however that these concepts may be hard for participants to judge independently. Indeed, finding objective physiological markers for these psychological constructs was the primary motivation of the present work.

## Data Availability Statement

Code for processing data will be made available at <u>parralab.org/resources</u>. Data cannot be publicly shared due to lack of IRB approval.

## Author Contributions

Ivan Iotzov

Roles: Conceptualization, data curation, formal analysis, investigation, methodology, software, visualization, writing - original draft, writing - review & editing

Lucas Parra

Roles: Conceptualization, funding acquisition, project administration, supervision, writing - original draft, writing - review & editing

## Acknowledgements

This work was supported in part by a grant from GN Resound. We thank Tanya Ignatenko for discussions leading up to this work and to Andrew Dittberner for posing the problem of how to measure hearing effort objectively.

